# Complemented palindrome small RNAs first discovered from SARS coronavirus

**DOI:** 10.1101/185876

**Authors:** Chang Liu, Ze Chen, Wenyuan Shen, Deshui Yu, Siyu Li, Yue Hu, Haishuo Ji, Wenjun Bu, Qingsong Wang, Shan Gao

## Abstract

In this study, we reported for the first time the existence of complemented palindrome small RNAs (cpsRNAs) and proposed cpsRNAs and palindrome small RNAs (psRNAs) as a novel class of small RNAs. The first discovered cpsRNA UCUUUAACAAG<u>CUUGUUAAAGA</u> from SARS coronavirus named SARS-CoV-cpsR-22 contained 22 nucleotides perfectly matching its reverse complementary sequence. Further sequence analysis supported that SARS-CoV-cpsR-22 originated from bat betacoronavirus. The results of RNAi experiments showed that one 19-nt segment of SARS-CoV-cpsR-22 significantly induced cell apoptosis. These results suggested that SARS-CoV-cpsR-22 could play a role in SARS-CoV infection or pathogenicity. The discovery of psRNAs and cpsRNAs paved the way to find new markers for pathogen detection and reveal the mechanisms in the infection or pathogenicity from a different point of view. The discovery of psRNAs and cpsRNAs also broaden the understanding of palindrome motifs in animal of plant genomes.

## Introduction

Small RNA sequencing (small RNA-seq or sRNA-seq) is used to acquire thousands of short RNA sequences with lengths of usually less than 50 bp. With sRNA-seq, many novel non-coding RNAs (ncRNAs) have been discovered. For example, two featured series of rRNA-derived RNA fragments (rRFs) constitute a novel class of small RNAs [1]. Small RNA-seq has also been used for virus detection in plants [2–4] and invertebrates [5]. In 2016, Wang *et al.* first used sRNA-seq data from the NCBI SRA database to prove that sRNA-seq can be used to detect and identify human viruses [6], but the detection results were not as good as those of plant or invertebrate viruses. To improve virus detection in mammals, our strategy was to detect and compare featured RNA fragments in plants, invertebrates and mammals using sRNA-seq data. In one previous study [7], we detected siRNA duplexes induced by plant viruses and analyzed these siRNA duplexes as an important class of featured RNA fragments. In this study, we detected siRNA duplexes induced by invertebrate and mammal viruses and unexpectedly discovered another important class of featured RNA fragments, which were complemented palindrome small RNAs (cpsRNAs). Among all the detected cpsRNAs, we found a typical 22-nt cpsRNA UCUUUAACAA<u>GCUUGUUAAAGA</u> from SARS coronavirus (SARS-CoV) strain MA15, which deserved further studies because mice infected with SARS-CoV MA15 died from an overwhelming viral infection with virally mediated destruction of pneumocytes and ciliated epithelial cells [8]. Although the palindromic motif TCTTTAACAAG<u>CTTGTTAAAGA</u> was already observed in a previous study [9], it never be considered to be transcribed as cpsRNAs before our studies.

The first discovered cpsRNA named SARS-CoV-cpsR-22 contained 22 nucleotides perfectly matching its reverse complementary sequence. In our previous study of mitochondrial genomes, we had reported for the first time a 20-nt palindrome small RNA (psRNA) named hsa-tiR-MDL1-20 [10]. The biological functions of hsa-tiR-MDL1-20 had been preliminarily studied in our previous study, while the biological functions of SARS-CoV-cpsR-22 were still unknown. In this study, we compared the features of siRNA duplexes induced by mammal viruses with those induced by plant and invertebrate viruses and found that siRNA duplexes induced by mammal viruses had significantly lower percentages of total sequenced reads and it seemed that they were only produced from a few sites on the virus genomes. One possible reason could be a large proportion of sRNA-seq data is from other small RNA fragments caused by the presence of a number of dsRNA-triggered nonspecific responses such as the type I interferon (IFN) synthesis [11]. Another possible reason could be the missing siRNA duplexes or siRNA fragments functions in cells by interaction with host RNAs or proteins. Based on this idea, we suspected that SARS-CoV-cpsR-22 could play a role in SARS-CoV infection or pathogenicity. Then, we performed RNAi experiments to test the cellular effects induced by SARS-CoV-cpsR-22 and its segments.

In this study, we reported for the first time the existence of cpsRNAs. Further sequence analysis supported that SARS-CoV-cpsR-22 could originate from bat betacoronaviruses. The results of RNAi experiments showed that one 19-nt segment of SARS-CoV-cpsR-22 significantly induced cell apoptosis. This study aims to provide useful information for a better understanding of psRNAs and cpsRNAs, which constitute a novel class of small RNAs. The discovery of psRNAs and cpsRNAs paved the way to find new markers for pathogen detection and reveal the mechanisms in the infection or pathogenicity from a different point of view.

## Results and Discussion

### Comparison of siRNA-duplexes induced by plant, invertebrate and mammal viruses

In this study, 11 invertebrate viruses were detected using 51 runs of sRNA-seq data (**Supplementary file 1**) and two mammal viruses (H1N1 and SARS-CoV) were detected using 12 runs of sRNA-seq data. In our previous study, six mammal viruses were detected using 36 runs of sRNA-seq data [6]. The detection of siRNA-duplexes by 11 invertebrate and eight mammal viruses was performed using a published program in our previous study [7]. Then, we compared the features of siRNA duplexes induced by invertebrate viruses (**Figure 1A**) with those induced by plant viruses (**Figure 1B**). The results showed that the duplex length was the principal factor to determine the read count in both plants and invertebrates. 21-nt siRNA duplexes were the most abundant duplexes in both plants and invertebrates, followed by 22-nt siRNA duplexes in plants but 20-nt siRNA duplexes in invertebrates. 21-nt siRNA duplexes with 2-nt overhangs were the most abundant 21-nt duplexes in plants, while 21-nt siRNA duplexes with 1-nt overhangs were the most abundant 21-nt duplexes in invertebrates but they had a very close read count to that of 21-nt siRNA duplexes with 2-nt overhangs. 18-nt, 19-nt, 20-nt and 22-nt siRNA duplexes in invertebrates had much higher percentages of total sequenced reads that those in plants. In addition, 18-nt and 19-nt siRNA duplexes had very close read counts and 20-nt and 22-nt siRNA duplexes had very close read counts in invertebrates. Since siRNA duplexes induced by mammal viruses had significantly lower percentages of total sequenced reads, the comparison of siRNA-duplex features between mammals and invertebrates/or plants could not provide meaningful results using the existing public data with standard sequencing depth. However, as an unexpected result from siRNA-duplex analysis, we discovered cpsRNAs from invertebrate and mammal viruses.

**Figure 1.**
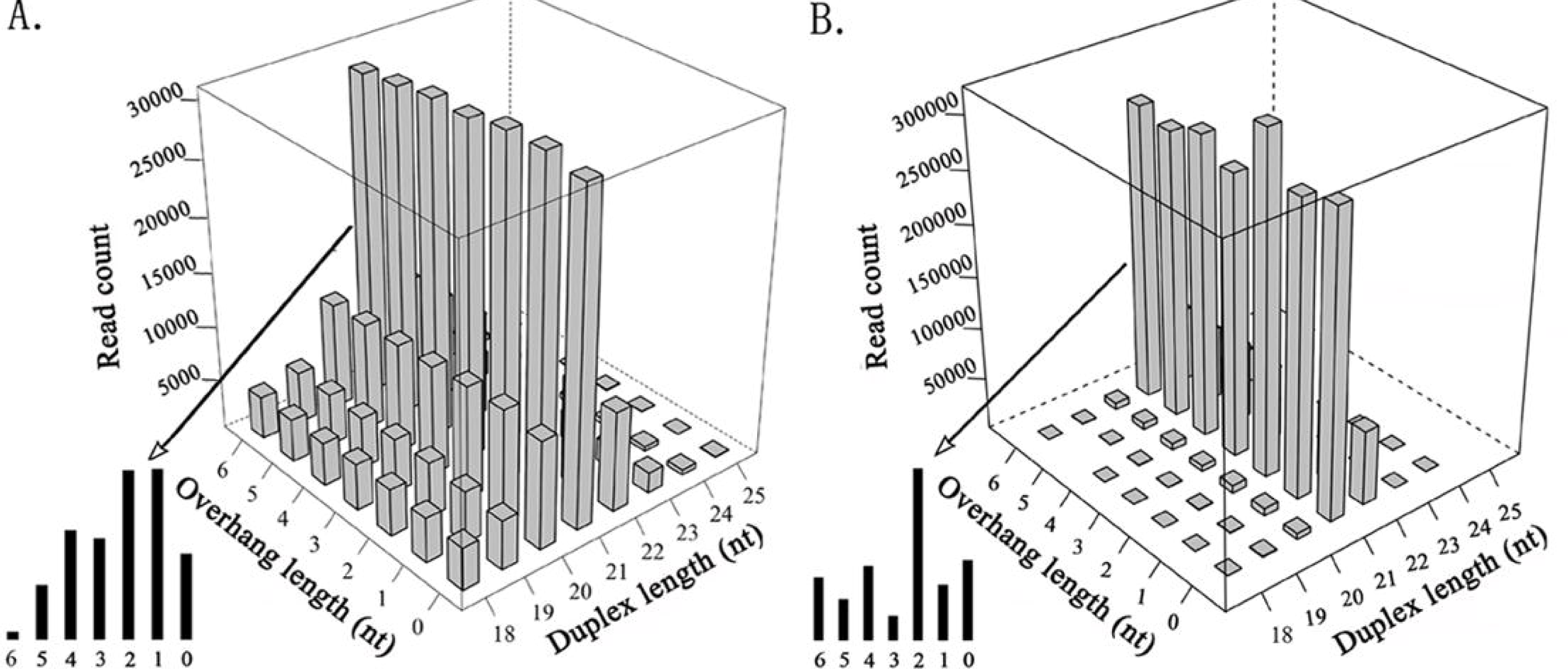
Comparison of siRNA-duplexes induced by plant and invertebrate viruses. All the cleaned sRNA-seq reads were aligned to viral genome sequences using the software bowtie v0.12.7 with one mismatch. The detection of siRNA duplexes was performed using the program duplexfinder [7]. **A.** The read count of siRNA duplexes varies with the duplex length and the overhang length, using data from 11 invertebrate viral genomes. **B.** The read count of siRNA duplexes varies with the duplex length and the overhang length, using data from seven plant viral genomes [7].

One typical cpsRNA UCUUUAACAAG<u>CUUGUUAAAGA</u> (DQ497008: 25962-25983) located in the orf3b gene on the SARS-CoV strain MA15 genome was detected in four runs of sRNA-seq data (SRA: SRR452404, SRR452406, SRR452408 and SRR452410). This cpsRNA was named SARS-CoV-cpsR-22, which contained 22 nucleotides perfectly matching its reverse complementary sequence (**Figure 2A**). We also detected one 18-nt and one 19-nt segment of SARS-CoV-cpsR-22, which could also be derived from siRNA duplexes (**Figure 2B**) but their strands (positive or negative) could not be determined. Among SARS-CoV-cpsR-22 and its two segments, the 19-nt segment was the most abundant and the 22-nt SARS-CoV-cpsR-22 was the least abundant.

**Figure 2.**
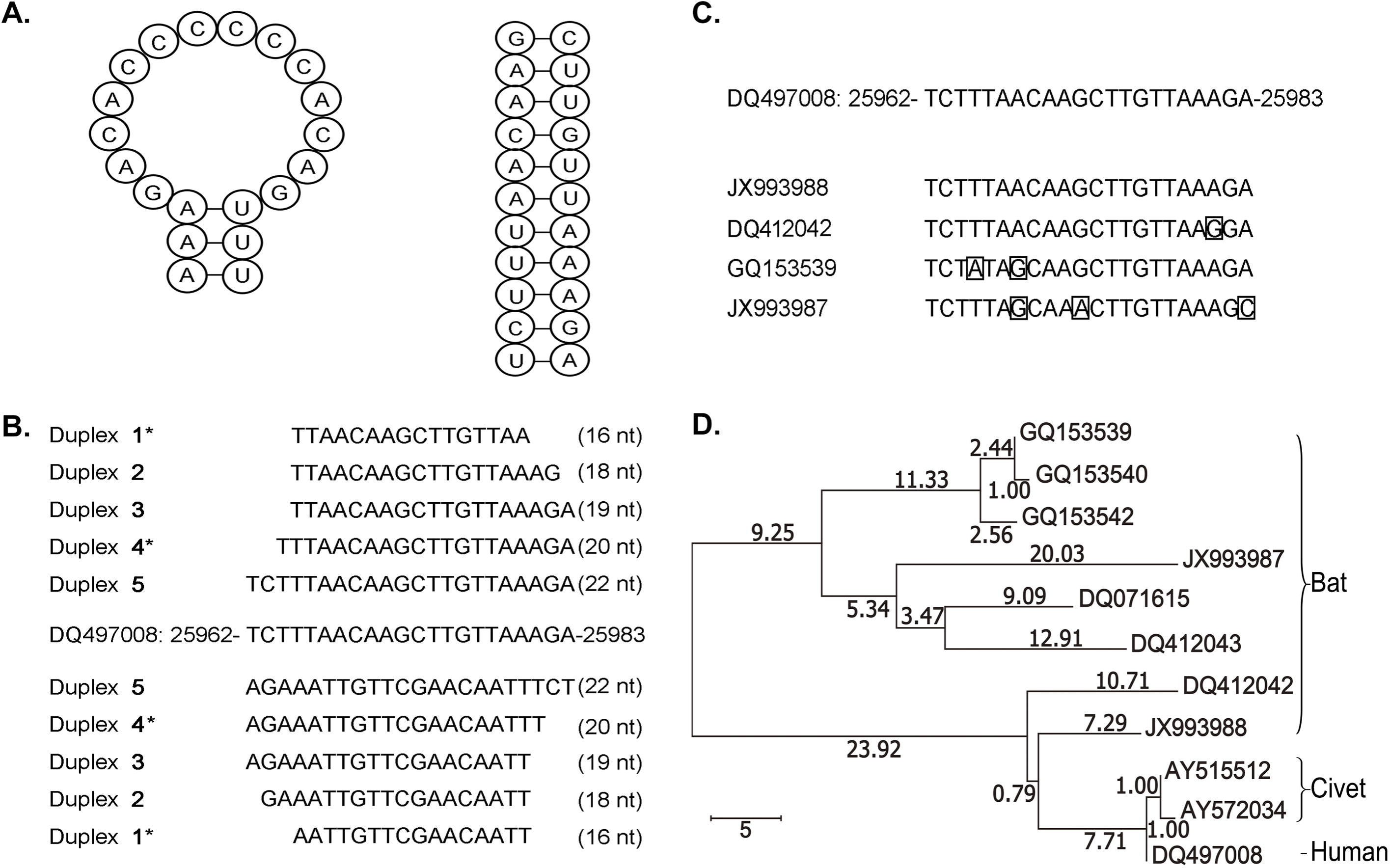
Clues to origins of SARS-CoV-cpsR-22. All the genome sequences are represented by their GenBank accession numbers (e.g. DQ497008). **A.** 16-nt, 18-nt, 19-nt, 20-nt and 22-nt siRNA duplexes were used for RNAi experiments. *16-nt and 20-nt had not been detected in this study. **B.** All the homologous sequences at the 22-nt locus on civet betacoronavirus genomes were identical to SARS-CoV-cpsR-22, while one of four genotypes at the 22-nt locus on bat betacoronavirus genomes was identical to SARS-CoV-cpsR-22. **C.** The psRNA (heteropalindromic) hsa-tiR-MDL1 and cpsRNA SARS-CoV-cpsR-22 can form hairpins. **D.** The phylogenetic tree was built by the Neighbor Joining (NJ) method using the orf3b gene from human betacoronavirus, eight homologous sequences from bat betacoronavirus and two homologous sequences from civet betacoronavirus. The branch’s length corresponds to an average number of nucleotide changes per 100 nucleotides.

### Discovery of psRNAs and cpsRNAs

Palindromic motifs are found in published genomes of most species and play important roles in biological processes. The well-known samples of palindromic DNA motifs include restriction enzyme sites, methylation sites and palindromic motifs in T cell receptors [12]. In this study, we found that palindromic or complemented palindrome small RNAs motifs existed ubiquitously in animal virus genomes, but not all of them were detected to be transcribed and processed into cpsRNAs probably due to inadequate sequencing depth of the sRNA-seq data. For example, we only found two psRNAs (CUACUGAC<u>CAGUCAUC</u> and AAGGUCUCC<u>CUCUGGAA</u>) from 14 palindrome motifs and one cpsRNA SARS-CoV-cpsR-22 from 29 complemented palindrome motifs with size from 14 to 26 nt (**Supplementary file 1**) on the SARS-CoV genome (GenBank: DQ497008.1) using four runs of sRNA-seq data (**Materials and Methods**). A DNA palindromic motif is defined as a nucleic acid sequence which is reverse complimentary to itself, while small RNAs which are reverse complimentary to themselves are defined as cpsRNAs. Accordingly, the typical psRNA should have a sequence which is 100% identical to its reverse sequence, but most psRNAs are heteropalindromic or semipalindromic as hsa-tiR-MDL1-20 reported in our previous study [10]. The psRNA hsa-tiR-MDL1-20 AAAGACACCCCCCACAGUUU (NC_012920: 561-580) contains a 14-nt region (underlined) which reads the same backward as forward and a 3-nt flanking sequence at the 5′ end which is reverse complimentary to a 3-nt flanking sequence at the 3′ end. With these 3-nt flanking sequences, hsa-tiR-MDL1-20 can form a hairpin as cpsRNAs usually do (**Figure 2A**). Although SARS-CoV-cpsR-22 is also typical, most of cpsRNAs have mismatches or InDels (Insertions/Deletions) in the reverse complimentary matches (**Supplementary file 1**). one example is a new Epstein-Barr Virus (EBV) microRNA precursor (pre-miRNA) with the length of 89 nt reported in our previous study [6]. This pre-miRNA sequence contains six mismatches and five insertions and only 87.64% (78/89) of the total nucleotides contributes to reverse complimentary matches.

### Clues to origins of SARS-CoV-cpsR-22

The previously unknown SARS virus generated widespread panic in 2002 and 2003 when it caused 774 deaths and more than 8000 cases of illness. Scientists immediately suspected that civet cats which were only distantly relatives to house cats may had been SARS-CoV’s springboard to human [13]. Later, scientists concluded that civets were not the original source of SARS. Further investigation showed that genetic diversity of coronaviruses in bats increased the possibility of variants crossing the species barrier and causing outbreaks of diseases in human populations [14]. To investigate the origins of SARS-CoV-cpsR-22, we obtained coronavirus genome sequences associated to bats, palm civets, rats and mice, monkeys, dogs, bovines, hedgehogs, giraffes, waterbucks and equines from the NCBI GenBank database. The results of sequence analysis showed that SARS-CoV-cpsR-22 was only located in the orf3b gene encoded by the betacoronavirus genome. Then, we blasted the orf3b gene from human betacoronavirus (GenBank: DQ497008.1) to all the obtained betacoronavirus genomes, except those for experiments on mice and monkeys. The results showed that the orf3b gene from human betacoronavirus had homologous genes from betacoronavirus in bats and palm civets (**Supplementary file 2**) rather than in other species. SARS-CoV-cpsR-22 also had homologous sequences at a 22-nt locus on bat and civet betacoronavirus genomes. All the homologous sequences at the 22-nt locus on civet betacoronavirus genomes were identical to SARS-CoV-cpsR-22, while one of four genotypes at the 22-nt locus on bat betacoronavirus genomes was identical to SARS-CoV-cpsR-22 (**Figure 2C**). Four genotypes at the 22-nt locus on bat betacoronavirus genomes had no, one, two and three mismatchs to SARS-CoV-cpsR-22 and their corresponding orf3b homologous sequences had identities of 96.77%, 96.13%, 87.96% and 85.16%. This suggested that one variant containing SARS-CoV-cpsR-22 could originate from betacoronavirus in bats, then be passed to palm civets and finally to human. This was consistent with the results of phylogenetic analysis using the orf3b homologous sequences from bat and civet betacoronavirus genomes (**Figure 2D**). In the phylogenetic tree, all human and civet betacoronavirus containing SARS-CoV-cpsR-22 was grouped into one clade. The nearest relative of the human and civet clade was the bat betacoronavirus (GenBank: JX993988.1) containing SARS-CoV-cpsR-22 and the next nearest relative was the bat betacoronavirus (GenBank: DQ412042.1) containing 22-nt homologous sequence with one mismatch to SARS-CoV-cpsR-22.

### Preliminary studies on biological functions of SARS-CoV-cpsR-22

Our previous study showed that the psRNA hsa-tiR-MDL1-20 contained the Transcription Initiation Site (TIS) and the Transcription Termination Site (TTS) of the human mitochondrial H-strand and could involve in mtDNA transcription regulation [10]. This inspired us to speculate that cpsRNAs could have similar biological functions and we investigated SARS-CoV-cpsR-22 using RNA interference (RNAi). Then, SARS-CoV-cpsR-22 and its 16-nt, 18-nt, 19-nt and 20-nt segments were designed as siRNA duplexes and introduced into PC-9 cells by pSIREN-RetroQ plasmid transfection (**Materials and Methods**). As a result, the 19-nt and 20-nt segment significantly induced cell apoptosis to 2.76 and 1.48 times at 48 hours after their transfection, respectively. Particularly, the 19-nt segment significantly induced cell apoptosis to 7.94 (36.04/4.54) times at 72 hours after their transfection (**Figure 3**). This corresponded to the 19-nt segment being the most abundant among all the segments of SARS-CoV-cpsR-22. Using the 19-nt segment, we also tested cell apoptosis in five other human cell lines and one mouse cell line. The results showed the 19-nt segment significantly induced cell apoptosis in the cell lines of A549, MCF-7 and H1299, but not in the cell lines of MB231, H520 and L929 (mouse). Since different cell types express specific genes, the 19-nt segment could silence cell-type specific transcripts to induce cell apoptosis through RNAi. These results suggested that the 19-nt segment had significant biological functions and could play a role in SARS-CoV infection or pathogenicity.

**Figure 3.**
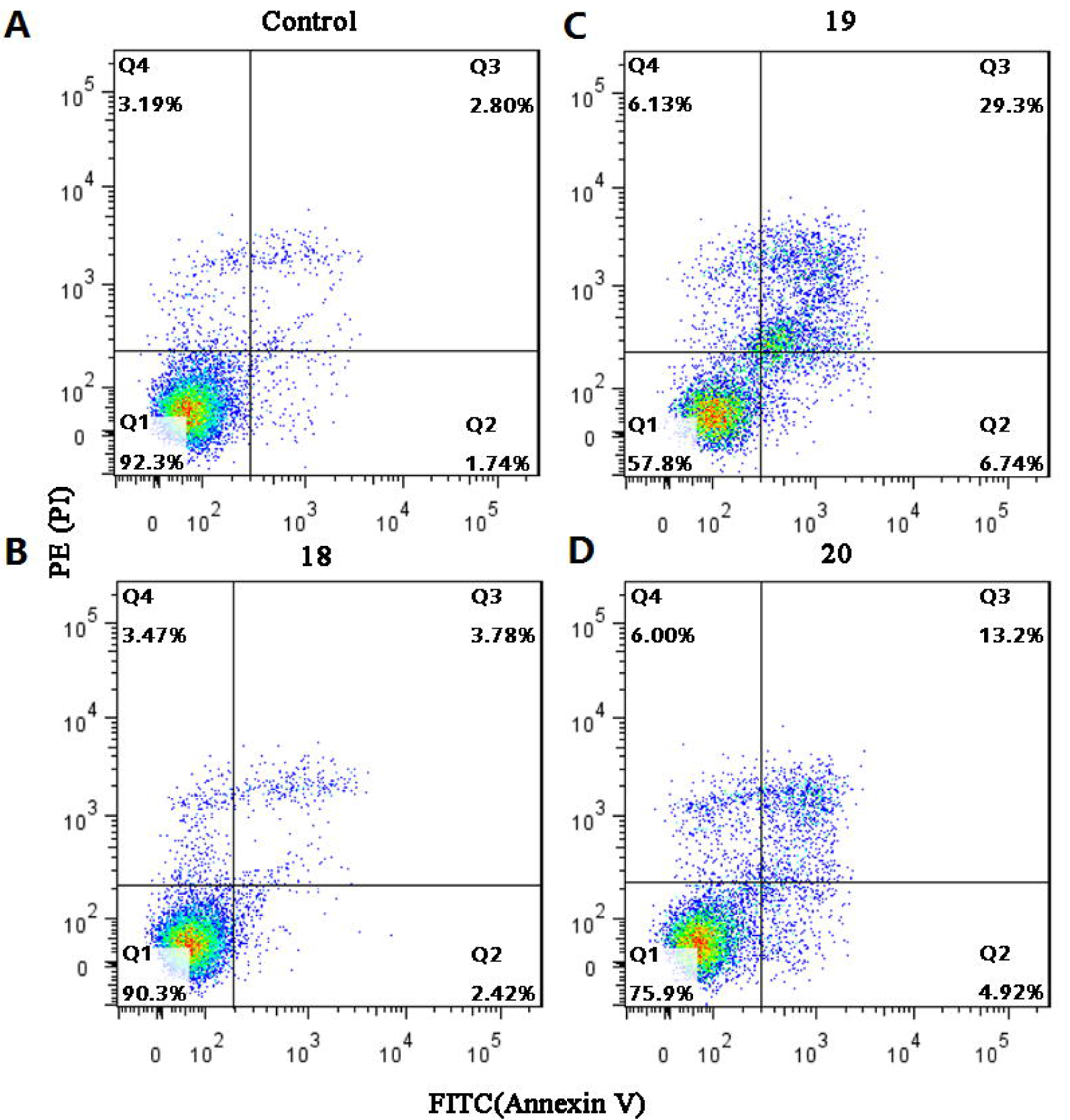
SARS-CoV-cpsR-22 induced cell apoptosis. PC-9 cells were divided into six groups named 22, 16, 18, 19, 20 and control for transfection using plasmids containing SARS-CoV-cpsR-22, its 16-nt, 18-nt, 19-nt, 20-nt segments. Each group had three replicate samples for plasmid transfection and cell apoptosis measurement. Each sample was processed following the same procedure. Cell apoptosis was measured at 72 h after transfection. The samples in this figure were selected randomly from their corresponding groups.

## Materials and Methods

### Datasets and data analysis

All sRNA-seq data were downloaded from the NCBI SRA database. The invertebrate and mammal viruses were detected from sRNA-seq data using VirusDetect [4] and their genome sequences were downloaed from the NCBI GenBank database. The description of sRNA-seq data and virus genomes is presented in **Supplementary file 1**. The cleaning and quality control of sRNA-seq data were performed using the pipeline Fastq_clean [15] that was optimized to clean the raw reads from Illumina platforms. Using the software bowtie v0.12.7 with one mismatch, we aligned all the cleaned sRNA-seq reads to viral genome sequences and obtained alignment results in SAM format for detection of siRNA duplexes using the program duplexfinder [7]. Statistical computation and plotting were performed using the software R v2.15.3 with the Bioconductor packages [16]. The orf3b gene from human betacoronavirus (GenBank: DQ497008.1), its 20 homologous sequences from bat betacoronavirus and nine homologous sequences from civet betacoronavirus were aligned using ClustalW2 (**Supplementary file 2**). After removal of identical sequences, the orf3b gene from human betacoronavirus, eight homologous sequences from bat betacoronavirus and two homologous sequences from civet betacoronavirus were used for phylogenetic analysis. Since these homologous sequences had high identities (from 85.16% to 99.78%) to the orf3b gene from DQ497008, the Neighbor Joining (NJ) method was used for phylogenetic analysis.

### RNAi and cellular experiments

Based on the shRNA design protocol [1], the sequences of SARS-CoV-cpsR-22, its 16-nt, 18-nt, 19-nt 20-nt segments (**Figure 2B**) and their control "CGTACGCGGAATACTTCGA" were selected to use as target sequences for pSIREN-RetroQ vector construction (Clontech, USA), respectively. PC-9 cells were divided into six groups named 22, 16, 18, 19, 20 and control for transfection using plasmids containing SARS-CoV-cpsR-22, its 16-nt, 18-nt, 19-nt, 20-nt segments and control sequences. Each group had three replicate samples for plasmid transfection and cell apoptosis measurement. Each sample was processed following the same procedure described below. At 12 h prior to transfection, the PC-9 cells were washed with PBS and trypsinized. Gbico RPMI-1640 medium was added into the cells, which were then centrifuged at 1000 rpm for 10 min at 4°C to remove the supernatant. Gbico RPMI-1640 medium (Thermo Fisher Scientific, USA) containing 10% fetal bovine serum was added to adjust the solvent to reach a volume of 2 μL and contain 2 × 10^5^ cells. These cells were seeded in one well of a 6-well plate for plasmid transfection. The transfection of 2 μg plasmids was performed using 5 μL Lipofectamine 2000 (Life technology, USA) following the manufacturer’s instructions. Cell apoptosis was measured with FITC Annexin V Apoptosis Detection Kit I (BD Biosciences, USA) following the procedure described below. The cells were washed by PBS, trypsinized and collectd using a 5-mL culture tube at 48 h or 72 h after transfection. The culture tube was then centrifuged at 1000 rpm for 10 min at 4°C to remove the supernatant. The cells were washed twice with cold PBS and resuspended in 1X Binding Buffer at a concentration of 1 × 10^6^ cells/mL. 100 μL of the solution (1 10^5^ cells) was transfered to a new culture tube with 5 μL of FITC-Annexin V and 5 μL PI. The cells were gently vortexed and incubated for 15 min at room temperature in the dark. 400 μL of 1X Binding Buffer was added to the tube. Finally, this sample was analyzed using a FACSCalibur flow cytometer (BD Biosciences, USA) within 1 hour. Apoptotic cells were quantified by summing the count of early apoptotic cells (FITC-Annexin V+/PI-) and that of late apoptotic cells (FITC-Annexin V+/PI+).

## Acknowledgments

We thank Yangguang Han and Jinke Huang from Tianjin KeYiJiaXin Technology Co., Ltd for his professional guidance on the RNAi and cellular experiments.

## Authors’ contributions

SG conceived this project. SG and QW supervised this project. SG, ZC and ZW analyzed the data. YS and HC curated the sequences and prepared all the figures, tables and additional files. CL, WS and YH performed qPCR experiments. SG drafted the main manuscript. DY and WB revised the manuscript. **All authors have read and approved the manuscript**.

